# Tuning Mitotic Recombination with Patterned DNA Nicks for Precision Mosaic Analysis

**DOI:** 10.1101/2025.10.30.685647

**Authors:** Yifan Shen, Ann T. Yeung, Bei Wang, Payton Ditchfield, Elizabeth Korn, Chun Han

## Abstract

CRISPR/Cas9-based mosaic analysis is a powerful tool for *in vivo* genetics but is limited by cytotoxicity and mutagenesis associated with DNA double-strand breaks (DSBs). Here, we establish Cas9-derived nickases as safer and more reliable alternatives for inducing mitotic recombination in *Drosophila*. We demonstrate that single-strand nicks are sufficient to generate mosaic clones and systematically dissect the parameters governing this process. We find that clone frequency can be controlled by the gRNA nicking pattern, with two distant nicks on the same DNA strand synergistically enhancing recombination by over nine-fold compared to a single nick. Based on these findings, we propose a mechanistic model for nick-induced crossover and provide a versatile toolkit for generating tissue-specific nickases. This work establishes nickase-based MAGIC as a superior method for high-fidelity clonal analysis, enabling more precise investigation of gene function in development and disease.

**SIGNIFICANCE STATEMENT:** The CRISPR/Cas9-based mosaic technique, MAGIC, is a versatile tool for *in vivo* biological investigations. However, its reliance on DNA double-strand breaks (DSBs) can cause significant, unintended cell damage. Here we establish that Cas9-derived nickases, which create gentler single-strand nicks, are a superior alternative. We show that nickases safely induce genetic mosaics in *Drosophila* by avoiding this cellular toxicity. By systematically dissecting the process, we discovered principles of gRNA design that allow clone frequencies to be ‘tuned’ for different experimental needs. This work provides a new mechanistic model for nick-induced genetic exchange, a high-fidelity “nickase-MAGIC” method, and a versatile toolkit for precision clonal analysis.

## INTRODUCTION

Mosaic analysis has been a cornerstone of developmental biology, enabling scientists to study the effects of lethal mutations, cell autonomy, cell lineage, and complex cell-cell interactions during tissue formation (1–3). The power of mosaic analysis lies in the ability to generate genetically distinct cell populations within a single organism, creating an *in vivo* environment where the consequences of homozygous mutations can be examined with high cellular resolution. For decades, mainstream mosaic techniques have relied on site-specific recombinases, such as Flippase (4, 5) and Cre (6, 7), to induce reciprocal exchange between homologous chromosomes in somatic tissues. However, these recombinases require specific recognition sequences that must be pre-engineered onto the test chromosome, limiting the application of these techniques to only a small subset of existing genetic resources and representing a significant bottleneck for studying novel mutations. We recently reported a recombinase-independent mosaic technique, mosaic analysis by gRNA-induced crossing-over (MAGIC) (8), which utilizes the CRISPR/Cas9 system to induce homologous recombination in somatic tissues. This approach eliminates the need for recombinase recognition sites and is thus compatible with virtually all existing wildtype (WT) and mutant chromosomes (8, 9), significantly expanding the applicability of mosaic analysis.

To induce homozygous clones, the MAGIC system uses Cas9 to generate DNA double-strand breaks (DSBs) at defined locations on a chromatid of a replicated chromosome. Using an intact chromatid of the homologous chromosome as a template, homology-directed repair (HDR) at the DSBs can catalyze crossovers between the two chromosomes, which can result in the generation of homozygous daughter cells following mitosis (8). However, DSBs are more commonly repaired by the alternative, error-prone non-homologous end joining (NHEJ) pathway. Competition for DSB repair by NHEJ is detrimental to MAGIC in at least three ways. First, the small insertions and deletions (indels) created by NHEJ can alter the guide RNA (gRNA) target sequence and thereby disable further CRISPR-mediated clone induction. Second, if the target sequences are not truly neutral, these same indel mutations may disrupt the function of nearby genes and lead to complicated and unexpected outcomes. Lastly, even if the target sequence is neutral, the simultaneous repair of multiple DSBs via end joining may result in large-scale chromosome aberrations, a risk well-documented in multiple systems (10). These competing repair outcomes collectively reduce the efficiency and reliability of Cas9-based mosaic analysis.

Cas9-derived nickases have been successfully used as Cas9 alternatives in a variety of CRISPR applications. These mutant variants of Cas9 have one of their two nuclease domains inactivated, restricting them to generating single-strand breaks (nicks) instead of DSBs (11, 12). The D10A variant has a defective RuvC domain and cleaves only the target strand complementary to the guide RNA. In contrast, the H840A variant, containing an inactive HNH domain, cleaves only the non-target strand containing the PAM sequence. These nickases have found increasingly more applications, from high-fidelity genome editing (12), allelic conversion (13), and gene drives (14) to prime editing (15). Because nickases do not generate DSBs, they are far less likely to cause the problems associated with mutagenic NHEJ repair mentioned above. However, a fundamental question remained: it was unknown whether nicks are sufficient to induce the crossovers required for a MAGIC experiment, given that DSBs are thought to be necessary for HDR. Furthermore, even if they can, how nickase-mediated clone generation would differ from that of Cas9 was unclear. Therefore, we sought to determine if nickases could be adapted for mosaic analysis, not only to circumvent the limitations of Cas9 but also to establish a new platform with higher fidelity and safety for *in vivo* genetic manipulation.

In this study, using *Drosophila*, we first confirm the deleterious effects of Cas9 in MAGIC experiments. We then show, surprisingly, that Cas9-derived nickases can induce mosaic clones efficiently and without these deleterious effects. Interestingly, Cas9 and nickases show different properties in MAGIC clone induction, influenced by both the target tissue and developmental timing. Using designer gRNAs, we uncovered the principles by which the DNA nicking pattern influences clone induction, indicating possible mechanisms in nick-induced crossing-over and allowing for greater control of clone frequency to match experimental purposes. Lastly, we developed a suite of tools for generating new tissue-specific, precursor-cell nickases. Our findings and reagents enable a more reliable and controllable next-generation of mosaic analysis through nickases, establishing a new standard for precision *in vivo* genetics.

## RESULTS

### Cas9, but not nickases, induces aberrant cells when paired with MAGIC gRNAs

In a standard MAGIC experiment (Figure S1A), to induce homozygous clones, Cas9 must generate a DSB at a targeted pericentromeric location on the chromosomal arm of interest during the G2 phase. HDR between the broken chromatid and an intact chromatid from the other homologous chromosome can cause a crossover between the two chromatids and subsequent exchange of the chromosomal segments distal to the crossover site. Clones homozygous for this chromosomal segment then result from the G2-X chromosomal segregation during the ensuing cell division (8). However, in addition to this ideal scenario, Cas9 may cut multiple chromatids simultaneously, and NHEJ may subsequently cause random chromosomal fusion and large-scale chromosomal aberrations (10), such as chromosomal non-disjunction and loss of chromosomal segments (Figure S1B and S1C). These types of chromosomal aberrations, predicted to be independent of MAGIC clone induction, would be detrimental to cells.

Indeed, in our applications of the MAGIC technique, we observed cytotoxicity and cell ablation consistent with the above predictions. For example, when we paired a ubiquitous Cas9 (*vas-Cas9*) and a previously reported MAGIC gRNA, *gRNA-40D2(Gal80)* (8), in a WT background, we observed low frequencies of dystrophy and ablation of *Drosophila* larval sensory neurons (Figure 1A-1E). The dystrophic class IV dendritic arborization (C4da) neurons (Figure 1B) exhibited greatly reduced dendritic arbors as compared to other normal-looking neurons (Figure 1A). Ablation was indicated by the absence of neurons innervating the larval epidermis (Figure 1C). Similarly, when *gRNA-40D2(Gal80)* was paired with *zk-Cas9* (8), which is expressed in the embryonic ectoderm, we observed many small and round epidermal cells in third instar larvae (Figures 1G and 1J). Such aberrant cells were absent in control animals (Figures 1F and 1J).

**Figure 1.**
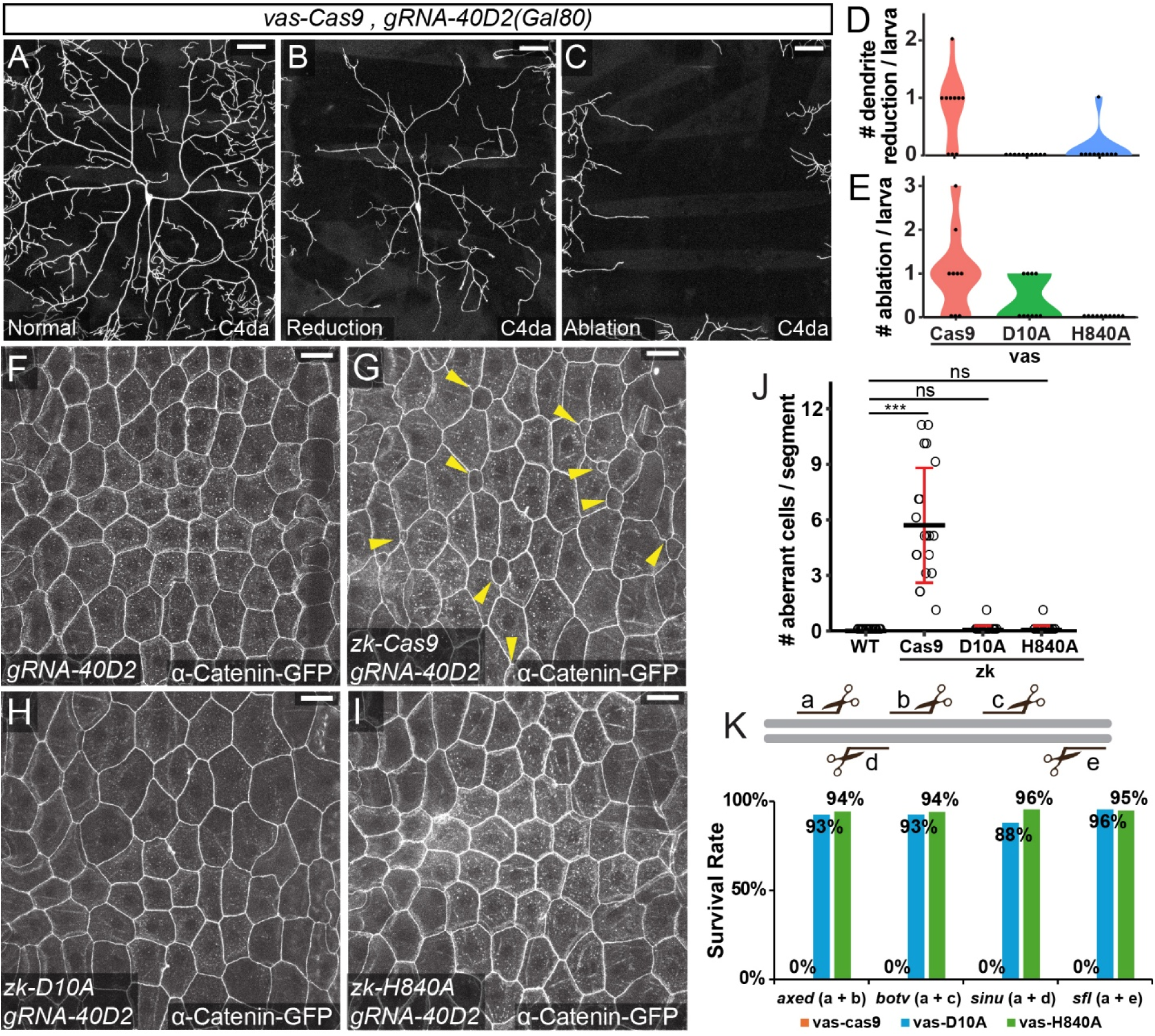
Cas9, but not nickases, induces aberrant cells when paired with MAGIC gRNAs. (A-C) Representative C4da neurons with normal dendritic field (A), reduced dendritic arbor (B), and neuronal ablation (C) in 3^rd^ instar larvae containing both *vas-Cas9* and *gRNA-40D2(Gal80)*. C4da neurons were labeled by *ppk-CD4-tdGFP*. (D and E) Violin plots of C4da neurons with dendritic reduction (D) and neuronal ablation (E). n = larvae numbers: *vas-Cas9* (n = 10), *vas-D10A* (n = 10), *vas-H840A* (n = 10). (F-I) Larval epidermis of *gRNA-40D2(Gal80)* alone (F), together with *zk-cas9* (G), *zk-D10A* (H), or *zk-H840A* (I). The epidermal cell shape was shown by the adherens junction marker α-Catenin-GFP. Yellow arrows point at the aberrant cells. (J) Numbers of aberrant cells in the larval epidermis. n = larval segments: WT (n = 20), *zk-Cas9* (n = 20), *zk-D10A* (n = 20), *zk-H840A* (n = 20). Black bar, mean; red bar, SD. one-way analysis of variance (ANOVA) and Tukey’s honest significant difference (HSD) test. ***p≤0.001, ns, not significance. (K) Survival rate of the progeny of crosses between gRNAs targeting essential genes and *vas-*driven Cas9 and nickases. The gRNA targeting pattern are shown on the top. Distance between gRNA target sites: *axed* (218 bp), *botv* (1476 bp), *sinu* (3 bp), and *sfl* (21,099 bp). Scale bar, 50 µm.

To ask if these aberrant and ablated cells are due to DSBs caused by Cas9, we tested *gRNA-40D2(Gal80)* with Cas9-derived nickases, which can only nick DNA but are otherwise similar to Cas9 (11). Indeed, nickases driven by the same *vas* enhancer caused much fewer defects in C4da neurons (Figures 1D and 1E): In the tested samples, *vas-D10A* showed reduced neuronal ablation and no dendrite reduction, while *vas-H840A* showed very rare dendrite reduction and no ablation. Similarly, in epidermal cells, aberrant epidermal cells were nearly eliminated when nickases driven by the *zk* enhancer were tested (Figures 1H-1J).

Cas9 is mutagenic due to its ability to generate DSBs. Although nickases lack this ability, a pair of gRNAs targeting opposite strands of DNA may still generate staggered DSBs with nickases. In our MAGIC designs, the gRNA-marker always expresses a pair of gRNAs targeting closely linked regions. To understand how such designs may affect the mutagenicity of nickases, we compared *vas*-driven Cas9 and nickases in knocking out four essential genes, including *axed* (16)*, botv* (17)*, sinu* (18), and *sfl* (19). Each gene was targeted by two gRNAs, with varying distances between the target sites and the strands the gRNAs target (the same or the opposite). Consistent with the ubiquitous nature of *vas-Cas9* expression, all four gRNA pairs resulted in 100% lethality in embryos or larvae when crossed to *vas-Cas9* (Figure 1K), confirming the efficiency of these gRNAs. In contrast, all gRNAs gave rise to ∼90% viable adults emerging from pupae when crossed to *vas-nickases* (Figure 1K). These data suggest that nickases are largely non-mutagenic even when nicking both DNA strands.

### Nickases can induce somatic recombination in *Drosophila* tissues

Given that nickases tend not to cause cell dystrophy that could complicate MAGIC analysis of mutations, we wondered if nickases are capable of inducing somatic recombination and thus can serve as safer alternatives of Cas9 in MAGIC experiments. To answer this question, we conducted clonal analysis in multiple larval tissues using *gRNA-40D2(Gal80)*, *vas*- and *zk*-nickases, and appropriate combinations of Gal4/UAS-fluorescent markers. With *vas-nickases*, we were able to readily detect clones in terminally differentiated cells, such as peripheral sensory neurons (Figure 2A) and central neurons in the larval ventral nerve cord (VNC) (Figure 2C), and proliferating tissues such as the wing imaginal disc (Figure 2B). With *zk-H840A*, we observed clones in polyploid epidermal cells (Figure 2D), indicating that nickases can induce crossovers when epidermal precursor cells are still dividing. These results collectively demonstrate the feasibility of using nickases to induce MAGIC clones across diverse *Drosophila* tissues.

**Figure 2.**
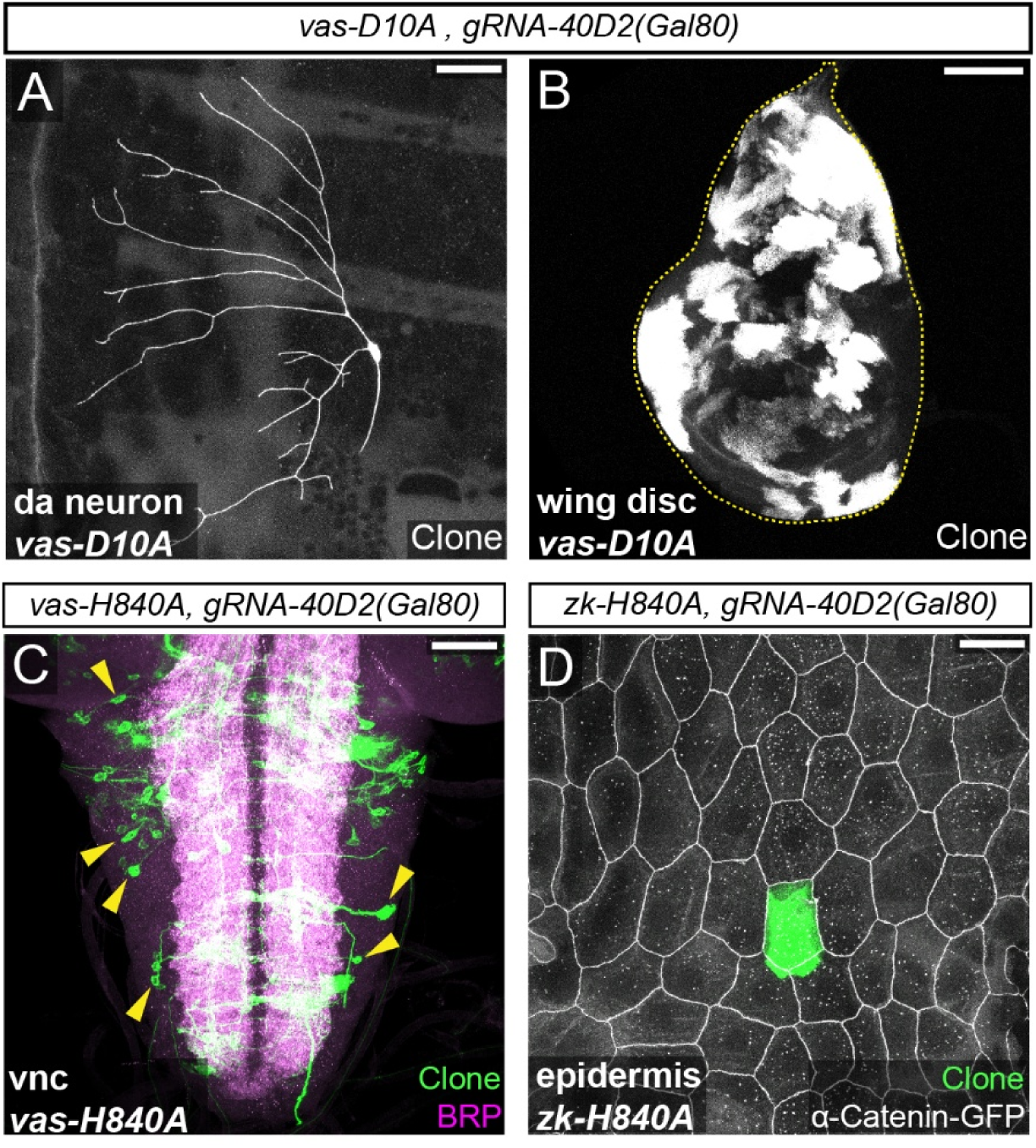
Nickases can induce somatic recombination in *Drosophila* tissues. (A) Class I da neuron clone on the larval body wall induced by *vas-D10A gRNA-40D2(Gal80)* and labeled by *RabX4-Gal4 UAS-MApHS*. (B) Clones in a larval wing disc induced by *vas-D10A gRNA-40D2(Gal80)* and labeled by *tub-Gal4 UAS-CD8-GFP*. (C) Neuronal clones in a larval ventral nerve cord induced by *vas-H840A gRNA-40D2(Gal80)* and labeled by *RabX4-Gal4 UAS-MapHS* (green). The neuropile is visualized by Bruchpilot (BRP) immunostaining (magenta). (D) A pMAGIC epidermal clone on the larval body wall induced by *zk-H840A gRNA-40D2(Gal80)* and labeled by *R38F11-Gal4 UAS-tdTom* (green). Epidermal junctions are labeled by *α-Catenin-GFP* (white). Scale bars: 100 µm in (B), 50 µm in (A), (C), and (D).

### Nickase expression pattern and gRNA targeting pattern affect MAGIC clone induction

To further explore the potential of nickases in MAGIC analysis, we examined three factors that may affect the ability of nickases to induce clones: the expression patterns of nickases, the identity of the nickase, and the gRNA nicking pattern.

We first compared Cas9 and nickases in the larval wing disc, a proliferating tissue, with *gRNA-40D2(Gal80)*. *vas*-driven Cas9 and nickases all showed efficient clone induction in the wing disc, and the differences among them in the sizes and numbers of clones were not statistically significant (Figures 3A and 3B). In contrast, *zk*-driven Cas9 and nickases showed drastic differences: While both *zk-nickases* were very efficient in generating clones, *zk-Cas9* resulted in very few but large clones in comparison (Figures 3A-3E). This difference between Cas9 and nickases could be due to the developmental timing of the *zk* enhancer activity. *zk* consists of a ventral repression element from the *zen* regulatory sequence (20) and two copies of the *Kruppel* CD1 enhancer element (21) and drives high expression in the dorsal ectoderm of the late blastoderm embryo (before stage 6) (21). Because the primordium of the wing disc is formed by a few progenitor cells in the ectoderm much later (stages 12–13) (22), DSBs at gRNA target sites in blastoderm embryos could generate premature mutations early in the wing lineage and disable the MAGIC system. In comparison, nickases do not mutate the gRNA target sequences and could continue to act after the wing primordium is formed. In any case, these results show that nickases could perform similarly or differently from Cas9 depending on their expression patterns.

**Figure 3.**
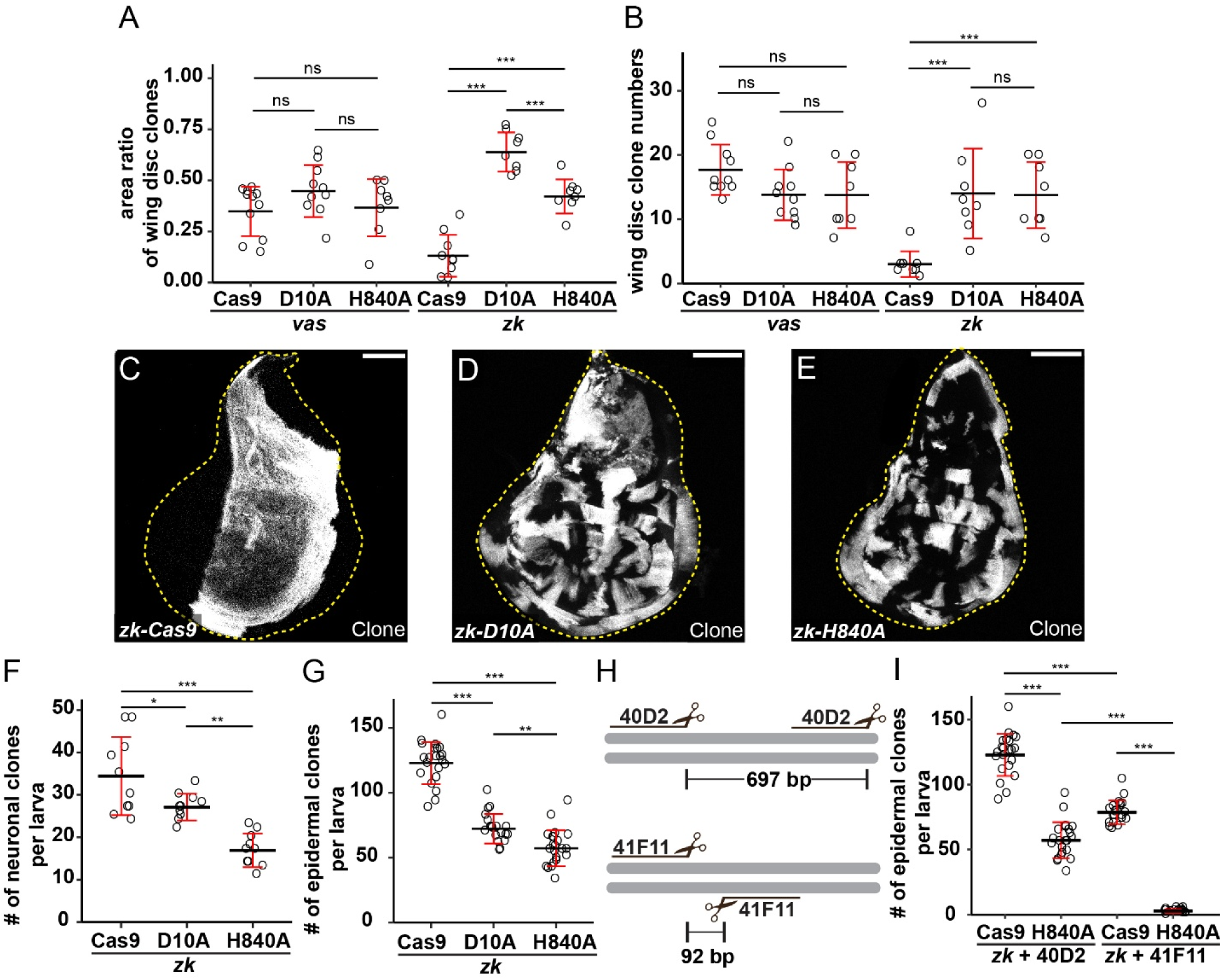
Nickase expression pattern and gRNA targeting pattern affect MAGIC clone induction. (A and B) Comparison of clone area ratios (A) and clone numbers (B) in larval wing discs among *vas*- and *zk*-driven Cas9 and nickases. Clones were induced by *gRNA-40D2(Gal80)* and labeled by *tub-Gal4 UAS-CD8-GFP*. n = disc numbers: *vas-Cas9* (n = 11), *vas-D10A* (n = 10), *vas-Cas9* (n = 8), *zk-Cas9* (n = 9), *zk-D10A* (n = 8), *zk-H840A* (n = 8). (C-E) Representative images of larval wing disc clones induced by *zk-Cas9* (C)*, zk-D10A* (D) and *zk-H840A* (E). Scale bar, 100 µm. (F and G) Frequencies of clones induced by *gRNA-40D2(Gal80)* and *zk*-driven Cas9 and nickases in larval peripheral neurons (F) and epidermal cells (G). For (F), n = larval numbers: *zk-Cas9* (n = 10), *zk-D10A* (n = 10), *zk-H840A* (n = 10). For (G), n = larvae number: *zk-Cas9* (n = 22), *zk-D10A* (n = 19), *zk-H840A* (n = 22). (H) Scheme of *gRNA-40D2* and *gRNA-41F11* target patterns. (I) Comparison of larval epidermis clone frequencies between two gRNAs. n = larvae number: *zk-Cas9 + 40D2* (n = 22), *zk-H840A + 40D2* (n = 22), *zk-Cas9 + 41F11* (n = 21), *zk-H840A + 41F11* (n = 18). In all plots, black bar, mean; red bar, SD. one-way analysis of variance (ANOVA) and Tukey’s honest significant difference (HSD) test. **p≤0.01, ***p≤0.001, ns, not significance.

In the above experiments, we also noticed that *zk-D10A* generated larger clones than *zk-H840A* in wing discs. To determine if D10A and H840A also perform differently in other tissues, we carried out clonal analysis in larval da neurons and epidermis with *zk-nickases* because they are more efficient than *vas*-driven nickases in these tissues. We found that D10A produced more clones than H840A in both tissues (Figures 3F and 3G). As a control, *zk-Cas9* induced more clones than both nickases (Figures 3F and 3G). These data suggest that, at least in the lineages of sensory neurons and epidermal cells where progenitor cells only exist during short developmental windows (23, 24), Cas9 outperforms nickases in inducing MAGIC clones. In addition, D10A is more effective than H840A in inducing clones.

To understand how the gRNA nicking pattern may affect the ability of nickases in inducing clones, we compared two existing pairs of MAGIC gRNAs targeting the second chromosome: 40D2 would generate two distant nicks on the same strand (697 bp apart), while 41F11 would make two closer nicks on opposite strands (92 bp apart) (Figure 3H). We tested clone frequency in the larval epidermis and found that, in both cases, *zk-Cas9* outperforms *zk-H840A*. Interestingly, compared to Cas9, H840A showed 53.4% drop in clone frequency with 40D2 but a much greater (96.6%) drop with 41F11 (Figure 3I). These data suggest that the nicking pattern can drastically affect clone induction by nickases. However, in these tests, because 41F11 is less efficient than 40D2 even with *zk-Cas9*, we could not separate the effects of the target locus and the nicking pattern.

### Designer gRNAs reveal principles of clone induction by nickases

To better understand the effect of the gRNA nicking pattern on nickase-induced crossover, we decided to compare gRNAs and their pairs targeting the same genetic locus but varying in the number, distance, and DNA strand of nicks, in the hope to minimize other variables. We chose the *Acp36DE* locus for picking gRNAs, because it encodes a seminal fluid-specific protein not required for *Drosophila* development (25) and thus can serve as a neutral but conserved gRNA target. *Acp36DE* also contains known single nucleotide polymorphisms (SNPs) in sequenced strains (26) that can disable specific gRNAs (Table S2), allowing us to test the impact of monoallelic vs biallelic cutting or nicking. In total, we picked 5 gRNA target sites that can give rise to 3 blunt-cut patterns with Cas9 and 5 nicking patterns with a nickase (Figure 4A). The 3 blunt-cut patterns include a single cut (a); two close cuts (a+b or a+d); and two distant cuts (a+c or a+e). The 5 nicking patterns include a single nick (a); two close nicks on the same strand (a+b); two distant nicks on the same strand (a+c); two close nicks on opposite strands (a+d); and two distant nicks on opposite strands (a+e) (Figure 4A). In these cases, close cuts/nicks are within 200 bp and distant cuts/nicks are more than 500 bp apart.

**Figure 4.**
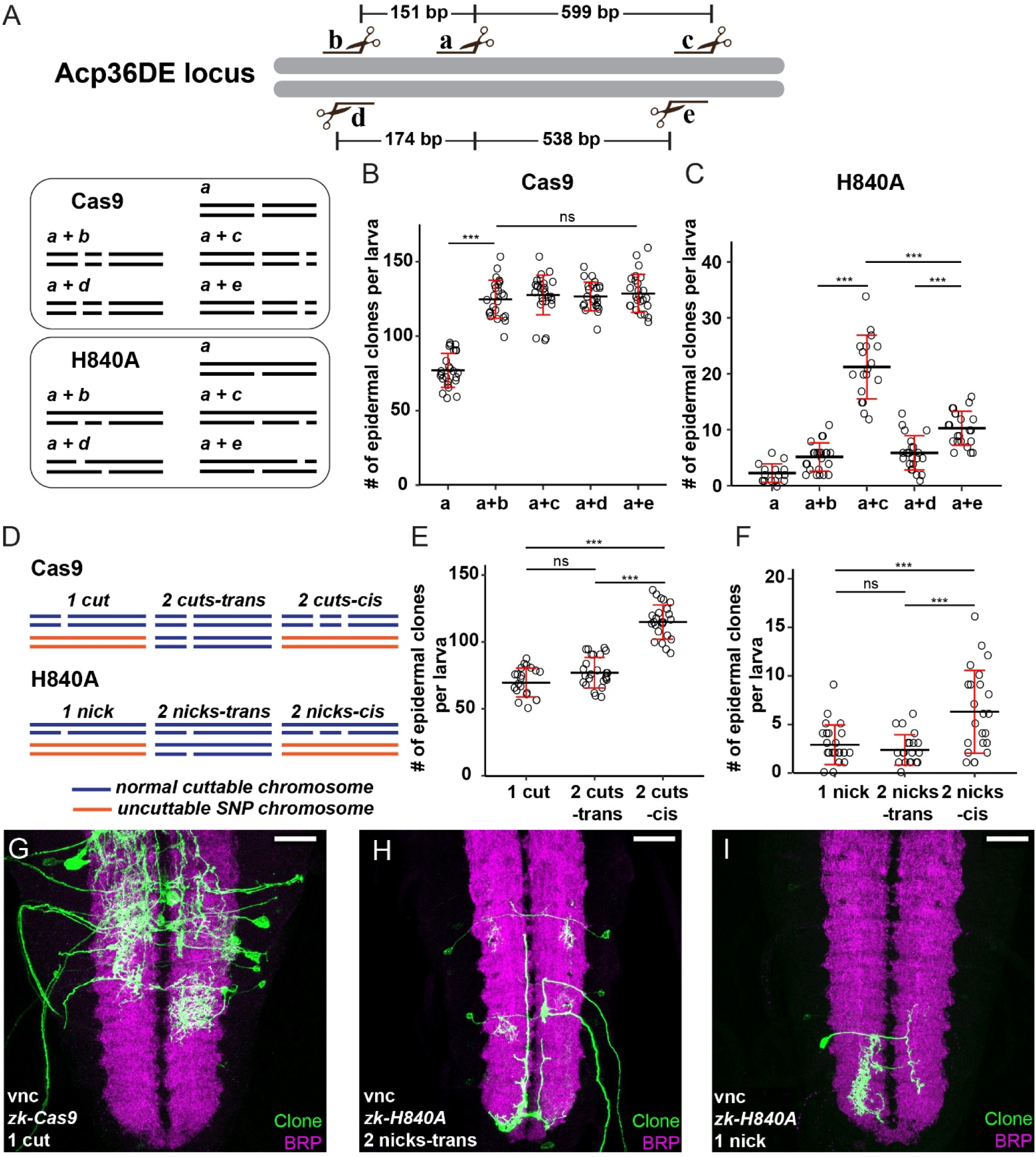
Designer gRNAs reveal principles of clone induction by nickases. (A) Scheme of gRNA target patterns at the *Acp36DE* locus and possible outcomes with Cas9 and H840A. (B) Clone numbers in the larval epidermis induced by *zk-Cas9*. n = larvae number: a (n = 25), a+b (n = 24), a+c (n = 26), a+d (n = 23), a+e (n = 25). (C) Clone numbers in the larval epidermis induced by *zk-H840A*. n = larvae number: a (n = 21), a+b (n = 23), a+c (n = 21), a+d (n = 22), a+e (n = 22). (D) Scheme of cutting/nicking patterns by designer gRNAs using uncuttable chromosomes. *gRNA-a* was used for 1 cut/nick (with chromosome 2 of DGRP 189) and 2 cuts/nicks-trans (with the *attP^VK00037^*chromosome). 2 cuts/nicks-cis utilized *gRNA-2-cis* and chromosome 2 of DGRP 189. The two target sites of *gRNA-2-cis* are 693 bp apart. (E and F) Clone numbers in the larval epidermis induced by *zk-Cas9* (E) and *zk-H840A* (F). n = larvae number: 1 cut (n = 21), 2 cuts-trans (n = 25), 2 cuts-cis (n = 25), 1 nick (n = 23), 2 nicks-trans (n = 21), 2 nicks-cis (n = 23). In all above experiments, epidermal clones were labeled by *R38F11-Gal4 UAS-tdTom.* (G-I) Representative image of neuronal clones labeled by *RabX4-Gal4 UAS-MApHS* in the larval ventral nerve cord using 1 cut (G), 2 nicks-trans (H), and 1 nick (I) patterns. In all plots, black bar, mean; red bar, SD. One-way ANOVA and Tukey’s HSD test. ***p≤0.001, ns, not significance. For figure G-I, scale bar 50 µm.

We first tested the efficiencies of these gRNAs by conducting clonal analysis in the larval epidermis with *zk-Cas9*. All four double cuts exhibited similarly efficient clone induction, and they gave rise to 1.62-1.67X higher clone frequency than the single cut (Figure 4B). These data confirm that all gRNAs are efficient and that dual gRNAs are more efficient in Cas9-mediated clone induction than single gRNAs. Importantly, the results also suggest that the distance between the two gRNA target sites is not a major factor determining the clone frequency.

We next tested these gRNAs with *zk-H840A*. Surprisingly, even a single nick was sufficient to induce clones, albeit at a very low frequency (Figure 4C). Adding another nick nearby (151 bp and 174 bp away) on either the same or the opposite strand increased the clone frequency by roughly two fold (2.16X and 2.46X, respectively) (Figure 4C), even though the frequency was still low. In contrast, adding a distant nick (538 bp and 599 bp away) caused much higher clone frequencies (Figure 4C), consistent with our initial tests of 40D2 and 41F11 (Figure 3I). Specifically, two distant nicks on opposite strands showed a 4.32 fold increase and those on the same strand showed a 9.08 fold increase. These data suggest that two distant nicks on the same strand synergistically increase DNA crossovers and are the most efficient configuration. Nevertheless, the highest efficiency with H840A is still 83% lower than that of the same gRNAs with Cas9.

Lastly, we tested whether cutting/nicking one or both homologous chromosomes would affect clone induction. Taking advantage of two chromosomes containing SNPs at the *Acp36DE* locus, we designed two new gRNAs that can target the locus on one chromosome but not the other. Using these gRNAs alone or together allowed us to create three scenarios for Cas9 (Figure 4D): one cut on a single chromosome (1 cut); two cuts at a single site on both homologous chromosomes (2 cuts-trans); and two cuts at two different sites separated by 693 bp on a single chromosome (2 cuts-cis). With a nickase, the same gRNA/chromosome combinations will similarly generate 1 nick, 2 nicks-trans, and 2 nicks-cis (Figure 4D). For both *zk-Cas9* and *zk-H840A*, 1 cut/nick and 2 cuts/nicks-trans generated similar clone numbers in the larval epidermis, while 2 cuts/nicks-cis exhibited much higher clone frequencies (Figures 4E and 4F). Across these experiments, Cas9 still outperformed H840A. These data suggest that whether only one or both homologous chromosomes are cut or nicked has minimal impact on crossing over, but two distant cutting/nicking sites on the same chromosome are much more efficient than one in inducing crossing over.

The 1 nick and 2 nicks-trans configurations in the above experiments are associated with the lowest clone frequencies. We reasoned that such low clonal frequencies could be useful to visualize neuronal morphology of individual clones in the densely packed central nervous system. Thus, we examined neuronal clones generated by 1 cut (as an example of a moderate clone frequency), 2 nicks-trans, and 1 nick in the VNC. As expected, 1 cut labeled many neurons with overlapping neuronal processes that are difficult to trace (Figure 4G). In comparison, both 2 nicks-trans (Figure 4H) and 1 nick (Figure 4I) sparsely labeled individual neurons, whose processes are spatially separated and readily traceable.

### Precursor-cell nickases can be generated in multiple ways

Our data above show that nickases are viable and more reliable alternatives of Cas9 in MAGIC experiments. However, currently very few nickases expressed in progenitor cells of specific tissues are available. To ease the generation of new tissue-specific nickases, we adopted two strategies that have been successfully used to make tissue-specific Cas9 strains (27, 28). First, for situations where a progenitor cell-specific enhancer is available, we developed two nickase Gateway destination vectors, pDEST-APIC-D10A and pDEST-APIC-H840A, using the pAPIC (attB P-element insulated CaSpeR) backbone optimized for enhancer-driven transgene expression (29) (Figure 5A). Tissue-specific enhancers can be conveniently swapped into this vector through the Gateway LR reaction to generate nickase-expression constructs. This cloning strategy is compatible with over 14,000 FlyLight (30) and VT (31) enhancers, whose expression profiles for multiple developmental stages and tissues in *Drosophila* are publicly available. *zk-D10A* and *zk-H840A* were generated using this method.

**Figure 5.**
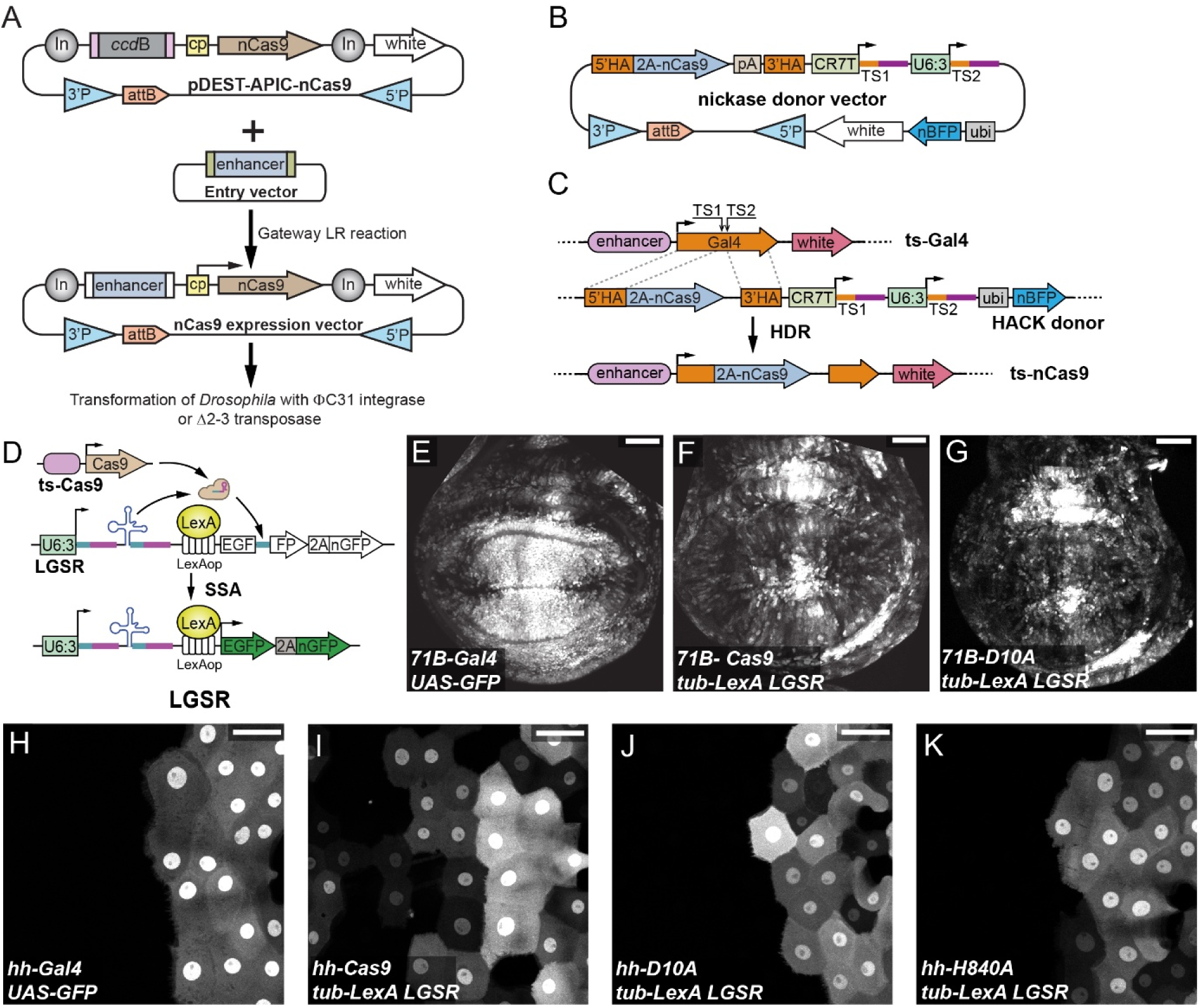
Precursor-cell nickases can be generated in multiple ways. (A) Diagram of Gateway cloning and transgenesis of nickase expression vectors. In, Gypsy insulator; cp, core promoter; 3′P and 5′P, P-element sequences; nCas9, nickase. (B) Diagrams of nickases HACK donor constructs. pA, polyA tail; TS, gRNA target sequence; P, P-element; HA, homology arm; ubi, a ubiquitous enhancer. pDEST-APIC-nCas9 and nickase donor vectors were constructed in pAC, a dual-transformation backbone (via PhiC31 and P-transposase) that carries a mini-white selection marker. (C) Diagram of Gal4-to-nickases conversion using a HACK donor line. The donor expresses two gRNAs (TS1 and TS2) targeting the tissue-specific (ts) Gal4, which results in in-frame incorporation of 2A-nickase into the Gal4 locus through HDR. The donor expresses ubi-nBFP that can be selected against when screening for convertants. (D) Diagram of LGSR. ts-Cas9, tissue-specific Cas9; SSA, single strand annealing; nGFP, nuclear GFP. (E-G) Activity patterns of *71B-Gal4* (E), 71B-Cas9 (F), and 71B-D10A (G) in the wing imaginal disc. (H-K) Activity patterns of *hh-Gal4* (H), *hh-Cas9* (I), *hh-D10A* (J), and *hh-H840A* (K) in the epidermis of the late 3^rd^ instar larva. In (E-K), Gal4 activity was visualized by *UAS-GFP.nls*, and Cas9/nickase activity was visualized by *tub-LexA LGSR*. For figure E-K, scale bar 50 µm.

Second, for situations where a Gal4 is known to be expressed in the progenitor cells of a particular tissue, we developed two nickase donor strains for converting the Gal4 into nickases through homology-assisted CRISPR knock-in (HACK) (Figure 5B) (32). Each nickase donor transgene carries a 2A-nickase coding sequence flanked by Gal4 homology arms, a gRNA cassette targeting the Gal4 coding sequence, and a ubiquitous nuclear BFP (nBFP) marker for distinguishing the donor chromosome (Fig. 5B). The nickase donors can be used to conduct knock-in experiments in the *Drosophila* germline entirely through genetic crosses, when combined with a germline cas9 and the Gal4 of interest. Successful in-frame insertion of a 2A-nickase in the Gal4 coding sequence will result in the expression of the nickase and a truncated and nonfunctional Gal4 in the original Gal4 expression pattern (Figure 5C).

In the HACK experiment, flies containing the converted nickase transgene can be identified using a Cas9 reporter. We previously developed GFP single-strand annealing reporter (GSR) to assist identification of Cas9 in HACK experiments (28). Driven by a ubiquitous promoter, however, GSR produces relatively low signals, which can pose challenges in animal screening. To enhance the signal, we generated LexAop-driven GSR (LGSR) (Figure 5D). Similar to GSR, LGSR expresses two gRNAs to target an interrupting sequence between two EGFP fragments in the transgene. In cells that express Cas9 or a nickase, the interrupting sequence is cut and the subsequent DNA repair by single-strand annealing between repeating sequences in the two EGFP fragments will reconstitute a complete EGFP-2A-nuclear GFP (nGFP) reading frame. However, different from GSR, LGSR can be expressed by a strong ubiquitous LexA, such as *tub-LexA*, to allow identification of larvae expressing GFP even in a small number of cells. As a proof-of-principle, we converted *hh-Gal4* and a wing disc-specific *71B-Gal4* (33) into Cas9 and/or nickase strains using *tub-LexA LGSR* as a screening reporter. These converted Cas9 and nickase strains exhibited similar activity patterns as the original Gal4 lines (Figures 5E-5K).

## DISCUSSION

In this study, we show that nickases can efficiently induce mitotic recombination in somatic tissues of *Drosophila* without the deleterious effects associated with Cas9, thus serving as much safer and more reliable alternatives in MAGIC experiments. By quantifying clone induction in several tissue types, we identified four factors that can influence the efficiency of clone induction by nickases: the expression pattern of the nickase, the target tissue, the nickase identity (D10A vs. H840A), and the gRNA nicking pattern. These results reveal potential mechanisms of nick-induced chromosome crossover and provide a clear framework for optimizing MAGIC design to match the experimental purposes. By establishing a toolkit for generating tissue-specific nickases, we extend robust and more reliable mosaic analysis to broader *Drosophila* tissues for addressing diverse biological questions.

### Possible mechanisms of nickase-induced crossover in somatic cells

Although a DSB is thought to be the canonical initiator of crossover during homologous recombination, we found that a single nick on a single homologous chromosome is sufficient to trigger mitotic recombination. This interesting observation is consistent with a recent study showing that DNA nicks in somatic tissues of *Drosophila* can induce HDR-based allele conversion (13). In that study, the nicked allele was repaired by copying the intact allele on the homologous chromosome. While it is unclear whether crossing-over accompanied the allele conversion, both processes could involve the same key intermediate of a D-loop generated during HDR (34, 35). Interestingly, early models of homologous recombination (HR) proposed nicks as the initiating lesions (36–38), although later experimental evidence strongly supported DSB-center models (39). So, how does a nick induce HDR? At least two mechanisms can explain this observation.

The most probable mechanism involves replication-coupled nick-to-DSB conversion in proliferating cells. DNA replication using a nicked strand as the template will result in a one-ended DSB (40, 41). At least in yeast cells, a one-ended DSB is efficiently converted into a double-ended DSB during S-phase (42), which is a potent substrate for HR. Alternatively, HDR could be induced by nick-to-gap processing. In this process, the nick is recognized by enzymes and widened into a single-stranded DNA (ssDNA) gap by exonucleases like EXO1 (43). The gap would be coated by the recombinase RAD51 to form a nucleoprotein filament (44), which can invade the intact homologous chromosome to initiate repair synthesis and form a D-loop (45). The D-loop is then resolved as a crossover, leading to mitotic recombination.

These two mechanisms can explain how the DNA nicking pattern affects nickase-mediated clone induction. Even though a single nick can be converted into a DSB when it encounters the replication fork, the nick is more likely to be repaired by direct ligation. Nick-to-DSB conversion in this way is stochastic and therefore relatively inefficient, which is why a single nick is the least effective configuration in inducing mitotic clones. We found that two nicks enhance mitotic recombination, but the degree of enhancement depends on which strand is nicked and the distance between the nicks. We hypothesize that opposite-strand and same-strand nicks may utilize different mechanisms. Opposite-strand nicks effectively create a staggered DSB independent of replication, increasing the chance of HDR. In contrast, two same-strand nicks are more likely processed into a ssDNA gap, which is also a substrate for HR and can similarly increase clone frequency. At a short distance (∼180 bp), both configurations double the clone frequency, suggesting that at this range, both lesion types are processed into substrates that promote HDR with similar efficiencies. Interestingly, two distant nicks (500-600 bp apart) further increase clone frequency, but same-strand nicks have a much greater effect than opposite-strand nicks. It is possible that simultaneous binding of two bulky nickases onto the same DNA is less efficient within a short range (e.g. 180 bp) due to steric hindrance, while at the 500-600 bp distance, the two nickases can bind DNA independently, without interfering with each other. Such lack of interference can explain the further increase of clone induction by opposite-strand nicks at the larger distance. In contrast, two same-strand nicks at this distance will create a long ssDNA gap, which provides a massive landing pad for RAD51 to form a long, stable filament. This filament structure should dramatically enhance the efficiency of the homology search and strand invasion required for crossover. In addition, this large ssDNA segment would almost guarantee that an incoming replication fork will stall and collapse, creating a DSB that robustly triggers crossover. Both mechanisms may contribute to the superior clone frequency of two distant same-strand nicks. These hypotheses can be tested in the future by examining nickase-induced clones when the relevant DNA repair pathways are manipulated.

### Effects of Cas9/nickase expression pattern and the proliferation property of the tissue

Our results also show that Cas9 and the two nickases have different efficiencies in inducing somatic clones, but their differences depend on the enhancer used to drive the nuclease expression and the type of the tissue examined. These differences can be explained by considering the nature of the lesions caused by the nucleases and the temporal window within which crossover must be induced.

A Cas9-generated DSB can efficiently activate HDR and induce crossover during S and G2 phases. In contrast, nicks require additional processing steps before HDR can be activated. Thus, Cas9 is expected to be more efficient than nickases in inducing crossover in a single cell cycle. However, DSBs can also be repaired by the error-prone NHEJ pathway throughout the cell cycle (34). If the gRNA target site is mutated during NHEJ-mediated repair, the target site is inactivated and will no longer be useful for crossover induction. In contrast, repair of nicks, either by direct ligation or HR, will restore the original sequence and make the target site available for the next round of nicking. Thus, while Cas9 is a potent but “single-shot” inducer, nickases can act iteratively, providing multiple opportunities for clone induction in tissues undergoing prolonged proliferation.

This prediction is reflected in our results in the wing imaginal disc, which is specified in mid-embryogenesis but maintains cell proliferation during the entire larval period. The peak expression driven by the *zk* enhancer is in the early embryo, when there are very few progenitor cells of the future wing disc. Cas9-induced DSBs at this early stage would either lead to mutations and inactivation of CRISPR, as reflected by the low clone frequency, or cause crossovers, as suggested by the observed large clones. In comparison, *zk*-nickases can keep generating nicks during the whole larval period and give rise to more, but smaller clones, even if the nickase level in this tissue is low post-embryogenesis. Unlike *zk*, the *vas* enhancer drives ubiquitous, and likely lower, expression throughout development (12). As long as the gRNA target sites are not mutated before the tissue starts to proliferate, *vas*-driven Cas9 and nickases would have enough time to induce clones.

The situation is different in epidermal cells and somatosensory neurons, whose precursor cells only exist for a short period before embryonic stage 13. Nickases thus have only very short temporal windows to induce crossover in these precursor cells. Supporting this idea, *zk-nickases* perform less efficiently than *zk-Cas9* in inducing clones in these tissues. We also observed very few clones in these tissues when using *vas-Cas9*, further supporting that *vas*-driven expression in early embryos is weak.

Lastly, we found that *zk-D10A* has a higher activity than *zk-H840A* in all three tissues examined. This observation is consistent with the recent finding that D10A has a higher cutting efficiency than H840A in *Drosophila* (14).

While our data strongly support a model where DNA repair and replication dynamics are the primary drivers of clone induction, we cannot rule out other contributing factors. For instance, the local chromatin environment at the target locus could influence the accessibility of the DNA to nickases or the recruitment of repair factors, potentially explaining some of the tissue-specific differences we observed. Future studies using gRNA target sites in different chromatin contexts could dissect these possibilities.

### Advantages and potentials of nickases in clonal analysis and precision genetics

While both Cas9 and nickases can be used in mosaic analysis, our results show that nickases have two clear advantages. The foremost is that nickases are much less mutagenic than Cas9 and rarely cause cell dystrophy. This improvement is crucial for accurately studying gene function and cell lineage without the confounding effects of cellular defects or cell death. Second, with nickases, it is possible to further adjust the clone frequency of MAGIC experiments over a larger dynamic range based on the principles revealed in this study, so that each experiment can be matched with a desired clone frequency. It is worth noting that a higher clone frequency is not always more desirable. For example, labeling a single axon within a densely packed neuropile may require a very low rate of clone induction. A gRNA that generates a single nick could be more appropriate for such purposes. As nickases are compatible with the existing MAGIC gRNA-markers kit (9) (https://bdsc.indiana.edu/stocks/misc/magic.html), one can pick a gRNA-marker based on the predicted nicking pattern to match the experimental interest. Alternatively, new gRNA-markers can be conveniently generated to induce a specific nicking pattern at a chosen locus (9), if existing ones do not fulfill the requirement.

To be useful for mosaic analysis, a nickase must be expressed in the precursor cells of the target tissue. The limited availability of precursor-cell nickases is currently a bottleneck for applying nickases in MAGIC experiments. To solve this problem, we developed a toolkit for making tissue-specific nickases through enhancer cloning or Gal4-to-nickase genetic conversion. New nickases created through these approaches can be characterized using the LGSR CRISPR reporter. Given the vast number of well-characterized enhancers and Gal4 lines available, it is feasible to generate precursor-cell nickases for most *Drosophila* cell types and tissues.

While demonstrated here in *Drosophila*, the fundamental DNA repair mechanisms we have leveraged are highly conserved across eukaryotes. Therefore, the principles of nickase-induced mosaic analysis established in this study may be readily transferable to other systems, including mammalian cell culture, organoids, and mice, opening new avenues for high-precision *in vivo* genetics in a broad range of biological contexts.

In conclusion, our findings on nickase-induced mosaic clones represent a significant advancement of the MAGIC technique. By offering a safer, more precise approach to clonal analysis while maintaining versatility across tissues, this improved MAGIC method has the potential to accelerate our understanding of gene function and developmental processes in *Drosophila* and other complex organisms.

## MATERIALS AND METHODS

### Fly Stocks and Husbandry

See the Key Resource Table (S1 Table) for details of fly stocks used in this study. Most fly lines were either generated in the Han lab or obtained from the Bloomington *Drosophila* Stock Center. *vas-Cas9, vas-D10A* and *vas-H840A* were kindly provided by Dr. Guichard and Dr. Bier. All flies were grown on standard yeast-glucose medium, in a 12:12 light/dark cycle, at 25°C. Virgin males and females for mating experiments were aged for 3-5 days.

MAGIC clones were labeled by *RabX4-Gal4 UAS-CD4-IFP2.0-T2A-HO1* or *UAS-MApHS* (46) in larval peripheral and central neurons, by *tub-Gal4 UAS-mCD8-GFP* in the larval wing discs, and by *R38F11-Gal4 UAS-tdTom* in the larval epidermis. In each MAGIC experiment, the appropriate gRNA(Gal80) chromosome was paired with a WT chromosome, so that the labeled clones contained homozygous WT chromosomes. The uncuttable chromosome illustrated in Figure 4D was derived from DGRP 189, which carries SNPs making the *Acp36DE* locus resistant to *gRNA-a* and *gRNA-cis*. The Cas9/nickase used in each experiment was indicated in figure legends.

To quantify the survival rate after knock-out of essential genes, *vas-cas9, vas-D10A*, or *vas-H840A* was crossed to gRNAs of essential genes. The survival rate was calculated as the number of adult progeny divided by the number of pupae.

### Molecular Cloning

#### D10A and H840A destination vectors

Mutagenesis cloning was performed by GenScript Biotech Corporation to replace the Cas9 coding sequence in the destination vector pDEST-APIC-Cas9 (Addgene 121657) (27) by D10A and H840A coding sequences, resulting in the corresponding nickase destination vectors pDEST-APIC-D10A and pDEST-APIC-H840A.

#### Nickase expression vector

*zk-D10A* and *zk-H840A* expression vectors were generated by Gateway LR reactions using the *zk* enhancer entry vector pENTR221-ZK2 (47) and the corresponding nickase destination vector.

#### gRNA expression vectors

gRNA target sequences were selected using the CRISPOR server (http://crispor.gi.ucsc.edu/). The gRNA target sequences for *botv* were cloned into pAC-U63-tgRNA-Rev as described (27). For *sinu,* gRNA target sequences were cloned into pAC-U63-QtgRNA2.1 as described (28). For *Acp36DE*, gRNA target sequences were cloned into pAC-U63-gRNA2.1-tubGal80(DE)-SV40 according to published protocols (9). The gRNA target sequences, the gRNA designs, and the cloning vectors are listed in Table S2.

#### Gal4-to-nickase HACK donor vectors

Mutagenesis cloning was performed by GenScript Biotech Corporation to replace the Cas9 coding sequence in pHACK(Gal4)-DONR(T2A-Cas9) (Addgene 194768) (28) by D10A and H840A coding sequences, resulting in the final construct pHACK(Gal4)-DONR(T2A-D10A) and pHACK(Gal4)-DONR(T2A-840A).

#### LGSR

A fragment containing the 13xLexAop2 enhancer, *Hsp70* core promoter, and a synthetic intron was PCR-amplified from pDEST-APLO (Addgene 112805) (47). Another fragment containing an EGFP single-strand annealing cassette was PCR-amplified from pAC-GSR (28). Both fragments were assembled into MluI- and BglII-digested pAC-GSR through NEBuilder HiFi DNA assembly (New England Biolabs, Inc) to generate pAC-LGSR.

#### tub-LexA

The pENTR221-tubP entry vector (48) was combined with the destination vector pDEST-APIC-LexAGAD (47) (Addgene 112807) in an LR Gateway reaction, resulting in the expression vector ptub-LexAGAD.

Injections were carried out by Rainbow Transgenic Flies (Camarillo, CA 93012 USA) or Genetivision (Stafford, TX 77477) to transform flies through φC31 integrase-mediated integration into attP docker sites.

### Conversion of Gal4 to nickases

A germline-specific *nos-Cas9* on the 3^rd^ chromosome was combined with the appropriate donor transgene on the 2^nd^ chromosome and a Gal4 insertion into the same fly through two sequential crosses. Male founder flies containing all three components were crossed to the reporter line, *tub-LexA LGSR*. The progeny was screened at 3^rd^ instar larvae under a Nikon SMZ18 fluorescence stereomicroscope, and those showing the expected GFP expression patterns were recovered for development into adulthood. The resulting flies were crossed to proper balancer stocks to separate the converted nickase chromosome from other components (the reporter, donor, or *nos-Cas9*).

### Live Imaging

Live imaging of larval epidermal cells, sensory neurons was performed as previously described (Amy’s paper). Animals were collected at 96 (for late third larvae) or 120 (for wandering third instar larvae) hours after egg-laying (AEL) and mounted in glycerol on a slide with vacuum grease as a spacer. Animals were imaged using a Leica SP8 confocal microscope. For epidermis, images were taken at the dorsal midline of A2 and A3 segments with a 40X NA1.3 oil objective, pinhole size 2 airy units, and a z-step size of 1 µm. For dendritic arborization neurons, images were taken from A1 to A7 hemi-segments with a 20X NA0.8 oil objective, pinhole size 2.5 airy units, and a z-step size of 3.5 µm.

### Imaginal wing disc imaging

Larval dissections were performed as described previously (27). Briefly, wandering third instar larvae were dissected in a small petri dish filled with cold phosphate-buffered saline (PBS). The anterior half of the larva was inverted. Then trachea and gut were removed. The sample was then transferred to 4% formaldehyde in PBS and fixed for 20 minutes at room temperature. After washing with PBS, the imaginal discs were placed in SlowFade Diamond Antifade Mountant (Thermo Fisher Scientific) on a glass slide. A coverslip was lightly pressed on top. Imaginal discs were imaged using a Leica SP8 confocal microscope with a 20X NA0.8 oil objective.

### Immunohistochemistry

Larval brains were dissected and fixed similarly to wing discs. After fixation, the brains were rinsed and washed at room temperature in PBS with 0.2% Triton-X100 (PBST). The samples were then blocked in PBST with 5% normal donkey serum (NDS) for 1 hour. Brains were then stained for 2 hours at room temperature with mouse anti-Brp antibody nc82 (1:100 dilution) in the blocking solution. After additional rinsing and washing, the samples were incubated with donkey anti-mouse antibodies conjugated with Cy5 (1:400) for 2 hours at room temperature. The samples were then rinsed and washed again before mounting and imaging.

### Image Analysis and Quantification

Image analyses were conducted in Fiji/ImageJ. Counting of reduction and ablation of da neurons and aberrant cells number of larval epidermis were completed manually after images were taken. To quantify the number of epidermal clones, the clones were detected based on thresholds to generate masks inside of an ROI covering larval T1-A3 segments. Then clone numbers within the ROI was then measured using Analyze Particles in Fiji/ImageJ. To quantify clones in wing discs, clones were detected based on a fixed threshold to generate masks, and the total area of clones and clone numbers in each disc was then measured using Analyze Particles. Counting of neuronal clones was completed manually during the imaging process. To count peripheral sensory neuronal clones on the larval body wall, third instar larvae were mounted laterally on slides, and the clones were counted in segments A1-A7 on one side of the larva under a Nikon SMZ18 stereomicroscope.

### Statistical Analysis

We first confirmed that the dependent variables were normally distributed using Shapiro-Wilk tests and that there was approximately equal variance across groups using Levene’s tests. One-way analysis of variance (ANOVA) with Tukey’s HSD test was then used for statistical comparison among groups. For additional information on the number of samples, see figure legends. R studio was used for all statistical analyses and all plots.

## ACKNOWLEDGMENTS

We thank Developmental Studies Hybridoma Bank (DSHB) for antibodies; Ethan Bier, Annabel Guichard, and Bloomington *Drosophila* Stock Center for fly stocks; Eric Alani for discussion; Norbert Perrimon, Tzumin Lee, Mariana Wolfner for advice; Mariana Wolfner, Eric Alani, Marcus Smolka, and Brooks Crickard for feedback on the manuscript. This work was supported by an NIH grant (R24OD031953) awarded to C.H..

---

**Figure S1.**
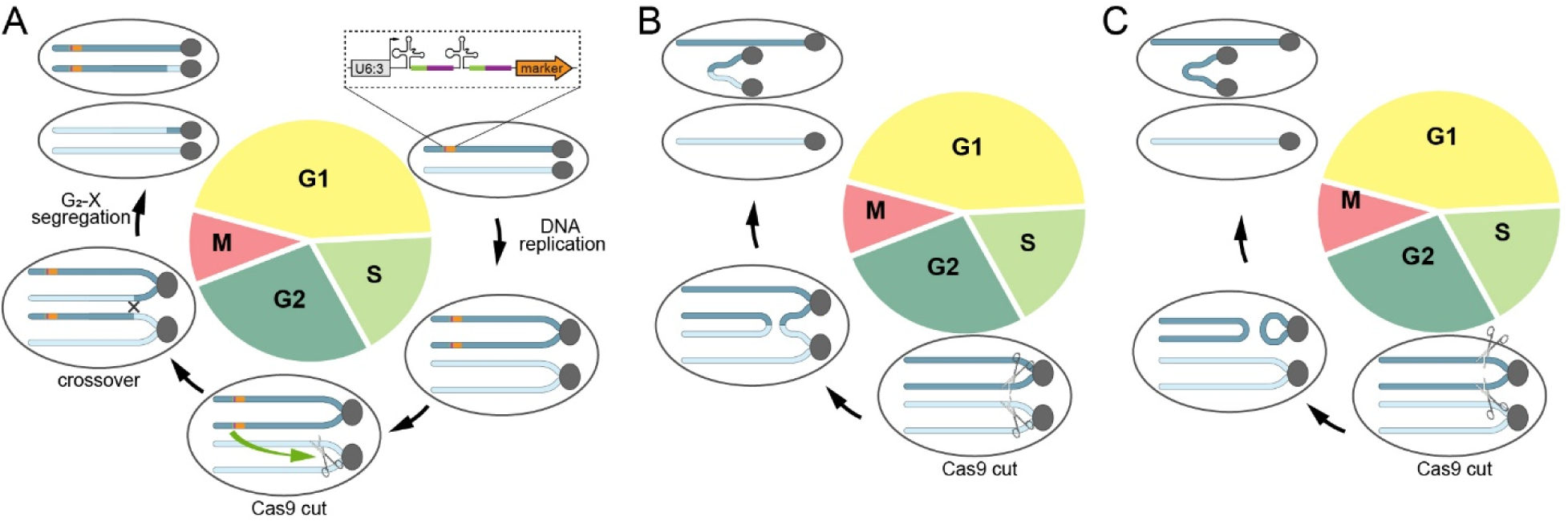
Principle of MAGIC and possible mechanisms of chromosomal aberrations. (A) Principle of MAGIC centered around the cell cycle. MAGIC utilizes a gRNA-marker transgene (enlarged box) inserted on a particular chromosomal arm. The gRNAs direct Cas9 to cut a pericentromeric site on the same arm. Ideally, Cas9 cuts a single chromatid during the G2 phase to induce crossover, which may lead to production of two daughter cells with one homozygous for the gRNA-marker and the other lacking the gRNA-marker. (B and C) Two possible scenarios in which multiple cuts of Cas9 lead to chromosomal fusion and non-disjunction in daughter cells. Other scenarios, such as chromosomal fusion at the G1 phase, are possible too.

## Notes

### Competing Interest Statement

The authors have declared no competing interest.

